# Molecular mechanism of substrate recognition by folate transporter SLC19A1

**DOI:** 10.1101/2022.09.09.507238

**Authors:** Yu Dang, Dong Zhou, Xiaojuan Du, Hongtu Zhao, Chia-Hsueh Lee, Jing Yang, Yijie Wang, Changdong Qin, Zhenxi Guo, Zhe Zhang

**Author notes:** Present address: Peking University First Hospital, Peking University Health Science Center, Beijing 100191, China (Xiaojuan Du).

## Abstract

Folate (vitamin B_9_) is the coenzyme involved in one-carbon transfer biochemical reactions essential for cell survival and proliferation, with its inadequacy causing developmental defects or severe diseases. Notably, mammalian cells lack the ability to *de novo* synthesize folate but instead rely on its intake from extracellular sources via specific transporters or receptors, among which SLC19A1 is the ubiquitously expressed one in tissues. However, the mechanism of substrate recognition by SLC19A1 has been unclear. Here we report the cryo-EM structures of human SLC19A1 and its complex with 5-methyltetrahydrofolate at 3.5-3.6 Å resolution and elucidate the critical residues for substrate recognition. In particular, we reveal that two variant residues among SLC19 subfamily members would designate the specificity for folate. Moreover, we identify intracellular thiamine pyrophosphate as the favorite coupled substrate for folate transport by SLC19A1. Together, this work has established the molecular basis of substrate recognition by this central folate transporter.

## Introduction

Folate (vitamin B_9_) is the coenzyme serving as the single-carbon donor in many biochemical reactions, e.g., the synthesis of purine and thymidylate, the metabolism of serine and methionine, and the methylation of nucleic acids and proteins^1^. Given such essential roles in cell growth, proliferation, and differentiation, folate inadequacy would lead to severe developmental defects or neurological disorders in humans^2,3^.

Mammalian cells lack the ability to *de novo* synthesize folate and must obtain it from extracellular sources such as foods. Three different systems are known for the transmembrane uptake of folate in mammals, i.e., the proton-coupled folate transporter SLC46A1^4^, the folate receptors (FRs)^5–7^, and the reduced folate carrier SLC19A1^8,9^. SLC46A1 is predominantly expressed in the gastrointestinal tract and is responsible for dietary folate absorption. Accordingly, SLC46A1 exhibits an optimal activity at acidic pH and couples folate transport to proton influx^10,11^. On the other hand, FRs take folate into cells via receptor-mediated endocytosis, primarily for folate delivery to the brain or folate retention in the kidney^7,12,13^. Notably, both SLC46A1 and FRs exert tissue-specific roles in folate transport. In contrast, SLC19A1 is ubiquitously expressed in the body and represents the major system of folate transport in diverse cell types^13^. For instance, though all three systems could facilitate the cellular uptake of antifolate drugs for cancer chemotherapy, SLC19A1 is the predominant route in many cancer cells^11,14,15^. Indeed, decreased expression or loss-of-function mutations of SLC19A1 in cancers would result in resistance to antifolate treatments^16^. Additionally, while SLC46A1 and FRs have equal affinities to folate and its reduced derivatives (e.g., 5-methyltetrahydrofolate, 5-MTHF), SLC19A1 shows a strong preference for the reduced derivatives^2^.

The structural mechanisms of folate transport by SLC46A1 and FRs have been elucidated^17–19^. However, despite its central role in folate uptake among different tissues, the molecular basis of substrate recognition by SLC19A1 has remained unclear. Here we report the cryo-EM structures of human SLC19A1 and its complex with 5-MTHF at 3.5-3.6 Å resolution and demonstrate the critical residues for substrate binding. In particular, we reveal two variant residues among SLC19 subfamily members, i.e., Arg133 and Gln377 in SLC19A1 *vs*. Glu138 and Met401 in SLC19A2, or Glu120 and Met384 in SLC19A3, being sufficient to designate the specificity for folate. Moreover, we identify intracellular thiamine pyrophosphate (TPP) as the favorite coupled substrate for folate transport by SLC19A1. These results have established the key mechanism of substrate recognition by SLC19A1.

## Results

### Structures of human SLC19A1 and its complex with 5-MTHF

Human SLC19A1 has 591 residues with a molecular weight of 65 kDa and is mainly composed of 12 transmembrane helices (TMs). As a result, the majority of the protein would be embedded in detergent micelles without obvious features when extracted from the cell membrane, limiting the structural determination by cryo-EM. To overcome this issue, we exploited the BRIL/Fab/Nb module^20–22^, which helped provide the apparent shape for particle alignment. The N-terminal 23 residues preceding the TM1 of SLC19A1 were replaced by the BRIL domain (Fig. 1a). Importantly, wild-type SLC19A1 or BRIL-SLC19A1 overexpressed in HEK293F cells exhibited comparable transport activity for a standard substrate [^3^H]-radiolabeled methotrexate ([^3^H]-MTX)^23^, indicating that the BRIL-tag would not affect the normal function of SLC19A1 (Fig. 1b and Supplementary Fig. S1a).

**Fig. 1.**
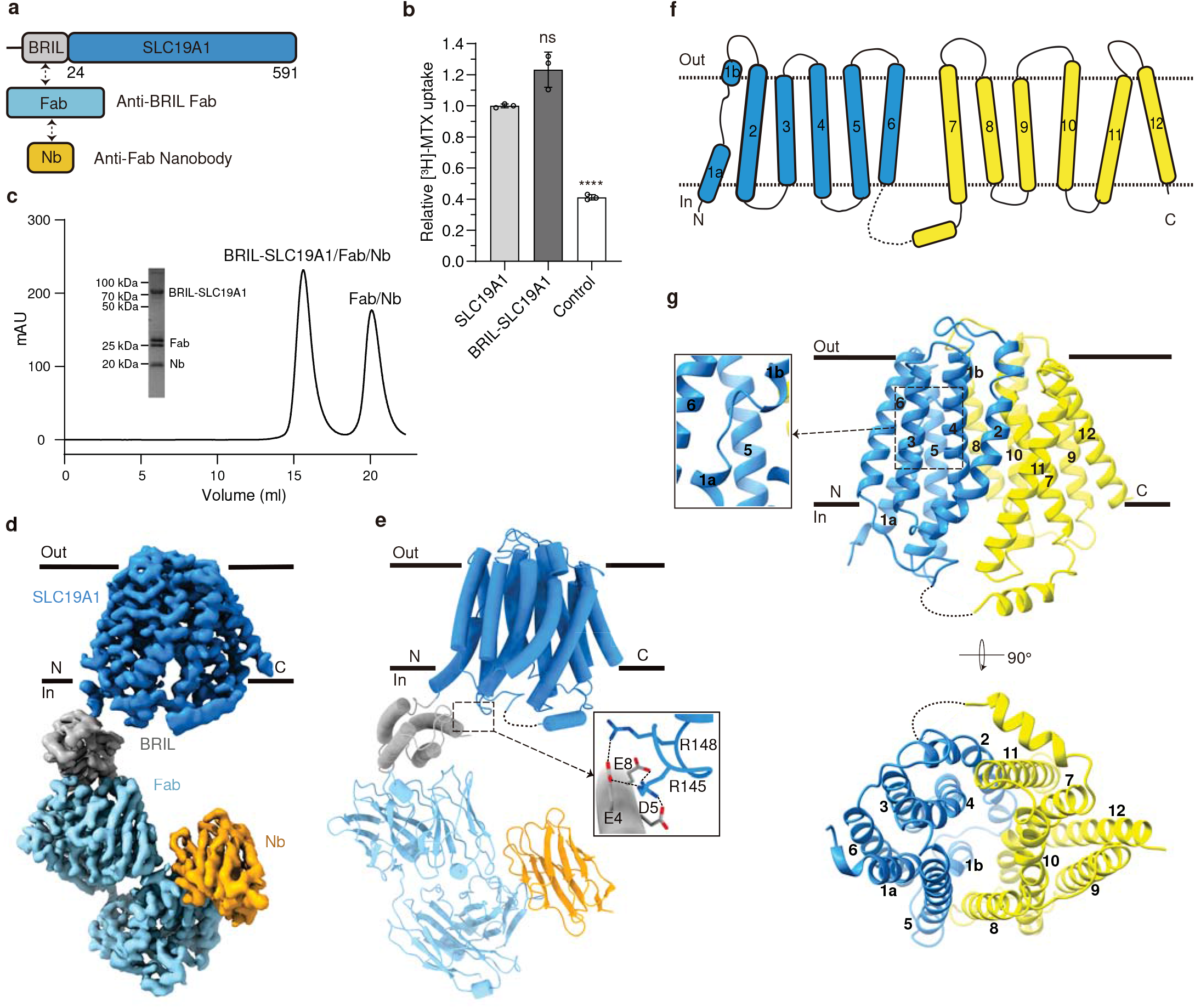
Cryo-EM structure of the BRIL-SLC19A1/Fab/Nb complex. **a** Schematic diagrams of BRIL-SLC19A1, anti-BRIL Fab, and anti-Fab Nanobody. **b** [^3^H]-MTX uptake assay to verify the function of BRIL-SLC19A1. The results are normalized to the activity of wild-type SLC19A1. All experiments were done in triplicates. (n = 3, mean ± s.d.). ns, non-significant; **** *P* < 0.0001 (T-test). **c** Profile of size exclusion chromatography (SEC) for the complex purification and the SDS-PAGE results to show the protein purity. **d** Cryo-EM map of the BRIL-SLC19A1/Fab/Nb complex. **e** Overall structure of the BRIL-SLC19A1/Fab/Nb complex. α-helices are shown in cylinders and β-strands are in ribbon. The residues of BRIL and SLC19A1 that are involved in electrostatic interactions are shown with side chains. **f** Cartoon diagram for the transmembrane (TM) domain of SLC19A1. The TM numbers are labeled, and the plasma membrane is indicated with dotted lines. **g** Ribbon presentation of the SLC19A1 structure in two views. Two half TM bundles are colored in blue (TM1-6) and yellow (TM7-12), respectively. A close-up view of the unwound region of TM1 is shown in an inset.

We purified the BRIL-SLC19A1 protein and then added anti-BRIL Fab and anti-Fab nanobody (Nb) to assemble the BRIL-SLC19A1/Fab/Nb ternary complex (Fig. 1c). The cryo-EM map of the ternary complex was collected and successfully reconstructed to 3.6-Å resolution (Fig. 1d, and Supplementary Fig. S2 and Table S1). In the structure of BRIL-SLC19A1, the last helix of BRIL rotated around its joint region with the TM1 of SLC19A1, and as a result, the four-helical bundle of BRIL resided parallel to the cell membrane. In addition, BRIL leaned on the intracellular loop between TM4 and TM5 of SLC19A1 via the electrostatic interactions between three acidic residues of BRIL (Glu4, Asp5, and Glu8) and two basic residues (Arg145 and Arg148) of SLC19A1 (Fig. 1e), stabilizing the current conformation of BRIL-SLC19A1. SLC19A1 adopted the classical major facilitator superfamily (MFS) fold^24,25^ with two discrete TM bundles (TM1-6 and TM7-12) (Fig. 1f, g), and all the TM regions were clearly resolved in the cryo-EM structure. SLC19A1 was present in the inward-facing conformation, i.e., the intracellular gate between TM4-5 and TM10-11 was open while the extracellular gate was closed by the regions of TM1, 2, and 7 (Fig. 1g). A notable feature was that a segment of TM1 (Ile41-Phe47) was unwound in the extracellular leaflet of the membrane (Fig. 1g). It has been documented that the discontinuity of transmembrane helices could play pivotal roles in transporters and ion channels by creating substrate-binding sites or providing flexible gating hinges^26–30^. On the other hand, the EM densities of the intracellular loop between TM6 and TM7 (residues 214-249) and the C-terminal cytoplasmic region (residues 452-591) of SLC19A1 were invisible (Fig. 1f, g), implicating their high motility and in line with their dispensable role for the transporter function^11^.

In parallel to the strategy of the BRIL module, we also identified one nanobody against SLC19A1 from a synthetic yeast-display library^31^. Using the nanobody-based legobody strategy^32^, we determined the cryo-EM structure of the SLC19A1/legobody complex to a medium resolution (~5 Å). The nanobody bound the TM6-7 linker region of SLC19A1, and the overall structure of the nanobody-bound SLC19A1 was almost identical to that observed in BRIL-SLC19A1 (Supplementary Fig. S3a). Given its higher resolution, the structure of BRIL-SLC19A1 (hereafter referred to as SLC19A1) was pursued in our further studies.

We next determined the cryo-EM structure of SLC19A1 in complex with its preferred substrate 5-MTHF at 3.5-Å resolution (Fig. 2a, and Supplementary Fig. S4 and Table S1). Similar to the apo-structure of SLC19A1, the SLC19A1/5-MTHF complex was in the inward-facing conformation (Fig. 2b). The 5-MTHF binding did not induce a significant conformational change of SLC19A1, as the root-mean-square deviation (RMSD) between the apo- and 5-MTHF-bound structures was 1.2 Å (Fig. 2c). Though the EM densities of the glutamate moiety of 5-MTHF were unresolved in the complex structure, the assignment of the substrate was unambiguously achieved. 5-MTHF resided inside the central cavity of SLC19A1 in the perpendicular position to the cell membrane, with the pterin ring pointing to the extracellular gate (Fig. 2b).

**Fig. 2.**
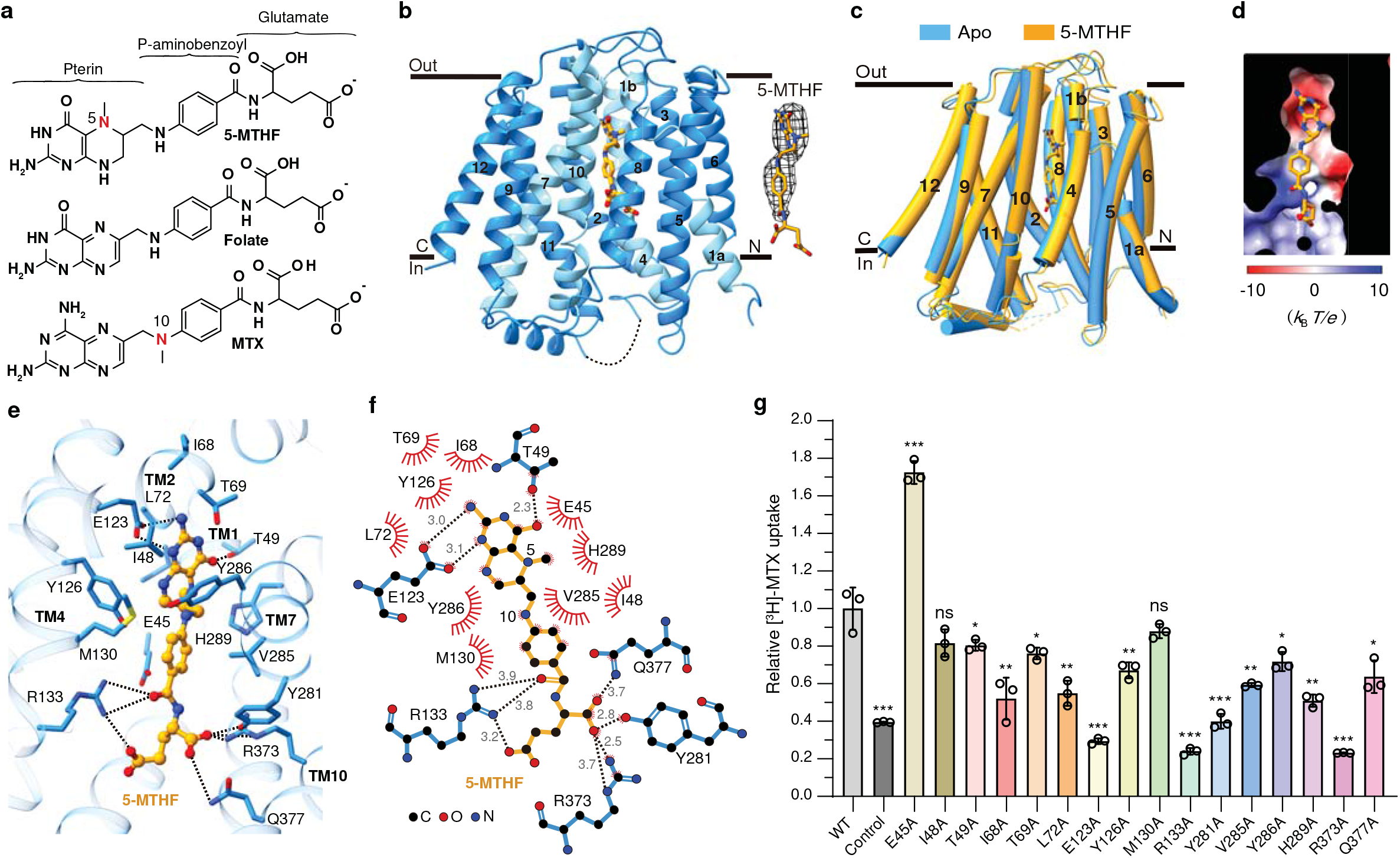
Cryo-EM structure of the SLC19A1/5-MTHF complex. **a** Chemical structures of several folate analogs, including 5-MTHF (5-methytetrahydrofolate), folate, and MTX (methotrexate). The constituent moieties of 5-MTHF are indicated. **b** Ribbon diagram of SLC19A1 in complex with 5-MTHF. The four TMs (TM1, 4, 7, and 10) interacting with 5-MTHF are colored in cyan and the other ones are colored in blue. 5-MTHF is shown in sticks and its cryo-EM densities are shown in black meshes. **c** Superposition of the apo and 5-MTHF-bound SLC19A1 structures. **d** The electrostatic potential (in units of *k*_B_*T*/*e*, where *k*_B_ is the Boltzmann constant, *T* is the absolute temperature and *e* is the elementary charge) of the substrate-binding pocket in SLC19A1, as calculated at pH 7.0 and 0.15 M concentrations of monovalent cations and anions. 5-MTHF is shown in sticks. **e** Ribbon presentation of the substrate-binding site in SLC19A1. The residues participating in 5-MTHF binding are indicated with side chains (< 4.0 Å). Hydrogen bonds and salt bridges are depicted as dashed lines. **f** A schematic summary of the interactions between SLC19A1 and 5-MTHF. The distances of the hydrogen bonds are indicated in angstroms. **g** The [^3^H]-MTX uptake activities of SLC19A1 mutants. The results are normalized to the activity of wild-type SLC19A1. All experiments were done in triplicates (n = 3, mean ± s.d.). ns, non-significant; * *P* < 0.01; ** *P* < 0.005; *** *P* < 0.001 (T-test).

### Mechanism of substrate recognition by human SLC19A1

In the structure of the SLC19A1/5-MTHF complex, the substrate-binding site was predominantly formed by TM1, 4, 7, and 10 of SLC19A1 (Fig. 2b). Notably, the electrostatic distribution of this binding site matched the charge characteristics of 5-MTHF (Fig. 2d), i.e., the polar pterin ring of 5-MTHF was wrapped in a negatively-charged pocket, and the glutamate moiety of 5-MTHF was located in a positively-charged environment.

We examined the critical residues involved in the substrate recognition of SLC19A1. On the extracellular side, the pterin ring of 5-MTHF formed hydrogen bonds with the side chains of Glu123 and Thr49 (Fig. 2e, f). At the same time, the pterin and *p*-aminobenzoyl groups of 5-MTHF were in close contact with an array of residues through van der Waals and hydrophobic interactions, including Glu45, Ile48, Ile68, Thr69, Leu72, Tyr126, Met130, Val285, Tyr286, and His289. It is worth noting that the extra methyl group on N5 nitrogen atom of 5-MTHF could enhance the hydrophobic interactions with SLC19A1 (Fig. 2a, e, and f), thus making 5-MTHF (*K*_t_ of 1-7 μM) a better substrate compared to folate (*K*_t_ of ~200 μM)^10^. On the intracellular side, the negatively-charged glutamate moiety of 5-MTHF was accommodated by two arginine residues (Arg133 and Arg373). Additionally, Tyr281 and Gln377 also participated in polar interactions with the glutamate moiety (Fig. 2e, f).

To verify the functional relevance of those residues involved in the substrate binding of SLC19A1, we mutated them individually and tested their effects on the transport activity in HEK293F cells (Fig. 2g and Supplementary Fig. S1b). Substitutions of Glu123, Arg133, Tyr281, and Arg373 with alanine abolished the transport activity of SLC19A1, consistent with their participation in the polar or electrostatic interactions with 5-MTHF. In comparison, the Q377A mutation only reduced the transport activity by ~40%, indicating that its polar interaction with 5-MTHF was less critical than that of the four residues above. In addition, T49A, I68A, T69A, L72A, Y126A, V285A, Y286A, or H289A mutation attenuated the transport activity by 20~50%, suggesting that their hydrophobic stacking with the pterin or *p*-aminobenzoyl ring would also contribute to substrate binding. Of importance, these key residues of SLC19A1 involved in the substrate recognition are mostly conserved among different species (Supplementary Fig. S5). In contrast, Ile48 and Met130 had minor roles as their mutations barely affect the function of SLC19A1. Intriguingly, mutation of Glu45 to alanine enhanced the transport activity by ~70%. This E45A mutation might better stabilize the unique loop structure of TM1 around the substrate-binding site (Supplementary Fig. S3b). In support of this notion, mutations of the residues adjacent to Glu45, e.g., G44R and S46I, have been identified in antifolate-drug resistant leukemia cells^33,34^. SLC19 subfamily contains three members, i.e., SLC19A1, SLC19A2, and SLC19A3.

Although sharing over 40% sequence identity, they engage different substrates, i.e., SLC19A1 transports folate, whereas SLC19A2 and SLC19A3 transport thiamine (vitamin B_1_)^10^. We investigated the mechanism designating the folate specificity of SLC19A1. By primary and ternary structural alignments, five out of the twelve residues of SLC19A1 comprising the binding site around the pterin and *p*-aminobenzoyl groups of 5-MTHF are not conserved in SLC19A2 and SLC19A3, i.e., Thr69, Leu72, Met130, Tyr286, and His289 (Fig. 3a-c). However, mutations of these residues to their cognates in SLC19A2 or SLC19A3 only slightly affected the transport activity of SLC19A1 (Fig. 3d and Supplementary Fig. S1c). We thus focused on the residues of SLC19A1 accommodating the negatively-charged glutamate moiety of 5-MTHF, i.e., Arg133, Tyr281, Arg373, and Gln377. While Tyr281 and Arg373 are conserved in SLC19A2 and SLC19A3, Arg133 and Gln377 become glutamate and methionine residues in SLC19A2 (Glu138 and Met401) and SLC19A3 (Glu120 and Met384), respectively (Fig. 3a-c). Replacing either of these two residues with their cognates in SLC19A2 and SLC19A3 (R133E or Q377M) completely abolished the function of SLC19A1 (Fig. 3d and Supplementary Fig. S1c). In addition, substituting the alanine residue adjacent to Arg133 to proline, i.e., A132P, caused the malfunction of SLC19A1^35,36^, elucidating the geometry restriction at Arg133. These results have suggested that these two variant residues among SLC19 subfamily members would be sufficient to determine the substrate specificity, with the negatively-charged glutamate and hydrophobic methionine residues of SLC19A2 and SLC19A3 precluding folate via the electrostatic or nonpolar repulsion on its glutamate moiety.

**Fig. 3.**
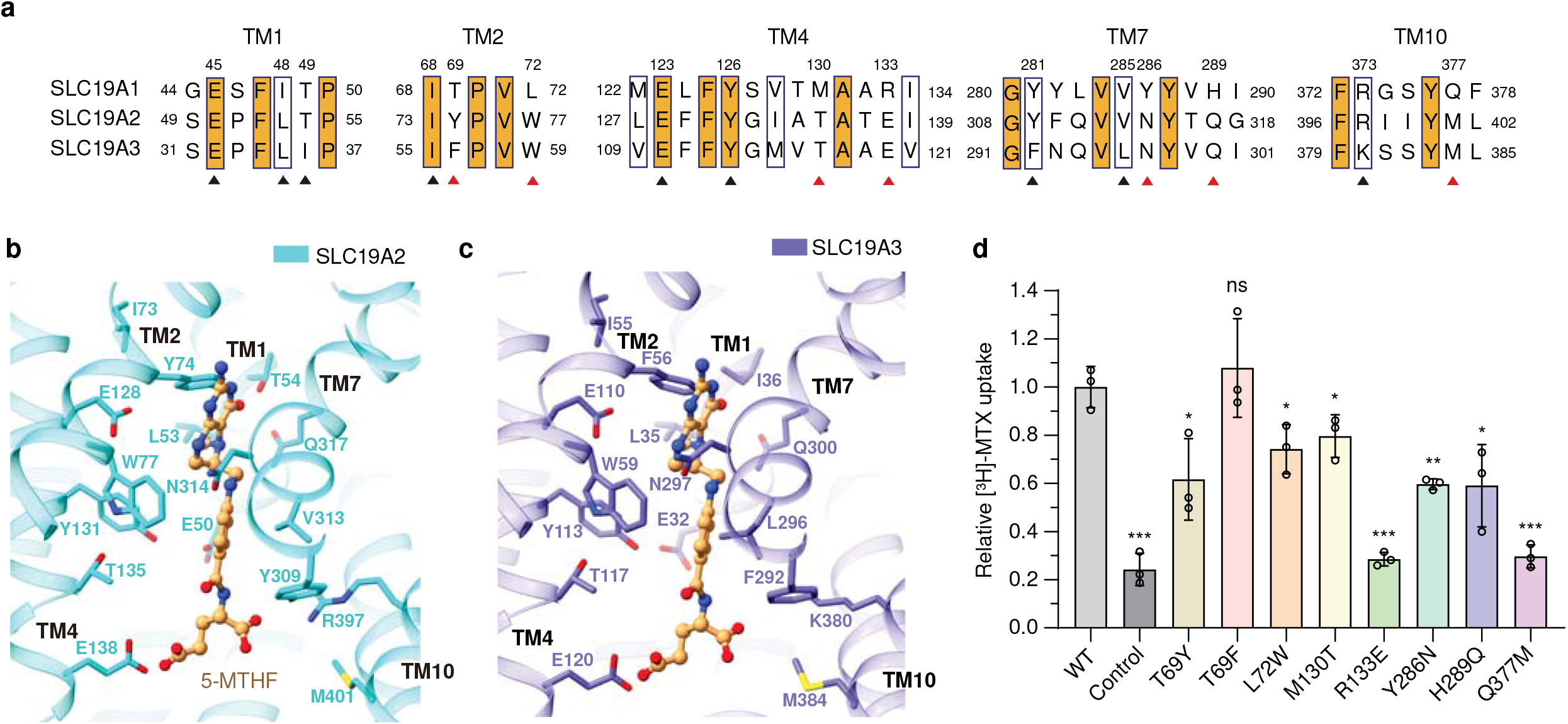
Analyses of the substrate discrimination mechanism of SLC19 subfamily members. **a** Sequence alignment of the three SLC19 family members. The partially conserved residues are indicated with blue boxes and the strictly conserved ones are further filled with orange color. The residues of SLC19A1 that involved in 5-MTHF binding are indicated. The conserved and non-conserved ones are denoted by black and red arrowheads, respectively. **b-c** Ribbon presentation of the 5-MTHF binding site in SLC19A2 and SLC19A3. The structures of SLC19A2 and SLC19A3 are predicted by AlphaFold. 5-MTHF is modeled into these structures based on the superposition with our SLC19A1/5-MTHF structure. The cognates of SLC19A2 and SLC19A3 corresponding to the 5-MTHF interaction residues of SLC19A1 are indicated. **d** Functional verification of the non-conserved residues for SLC19A1 using the [^3^H]-MTX uptake assay. The results are normalized to the activity of wild-type SLC19A1. All experiments were done in triplicates (n = 3, mean ± s.d.). ns, non-significant; * *P* < 0.01; ** *P* < 0.005; *** *P* < 0.001 (T-test).

### TPP is the favorite coupled substrate of SLC19A1

Although SLC19A2 and SLC19A3 both transport thiamine but not folate, most of the residues that comprise the substrate-binding pocket of SLC19A1, particularly those surrounding the pterin and *p*-aminobenzoyl groups of 5-MTHF, are highly conserved in SLC19A2 and SLC19A3 (Fig. 3a). Also, the non-conserved residues appeared to have a minor role in the transport function of SLC19A1 (Fig. 3d). These observations raised a tempting possibility that SLC19A1 might recognize specific types of thiamine derivatives as its substrate. Notably, SLC19A1 functions as an antiporter, i.e., coupling folate intake with the transport of another substrate in the opposite direction. In fact, a variety of organic phosphate anions were reported to be such coupled substrates of SLC19A1, including TPP, ATP (adenosine triphosphate), ADP (adenosine diphosphate), AMP (adenosine monophosphate), G6P (glucose 6-phosphate), and NAD^+^ (nicotinamide adenine dinucleotide)^23,37,38^. It thus came to our attention that the majority of thiamine would be metabolized in cells to its active form, the organic-phosphate derivative TPP (Fig. 4a), which is the coenzyme involved in biochemical reactions of decarboxylation^39^.

**Fig. 4.**
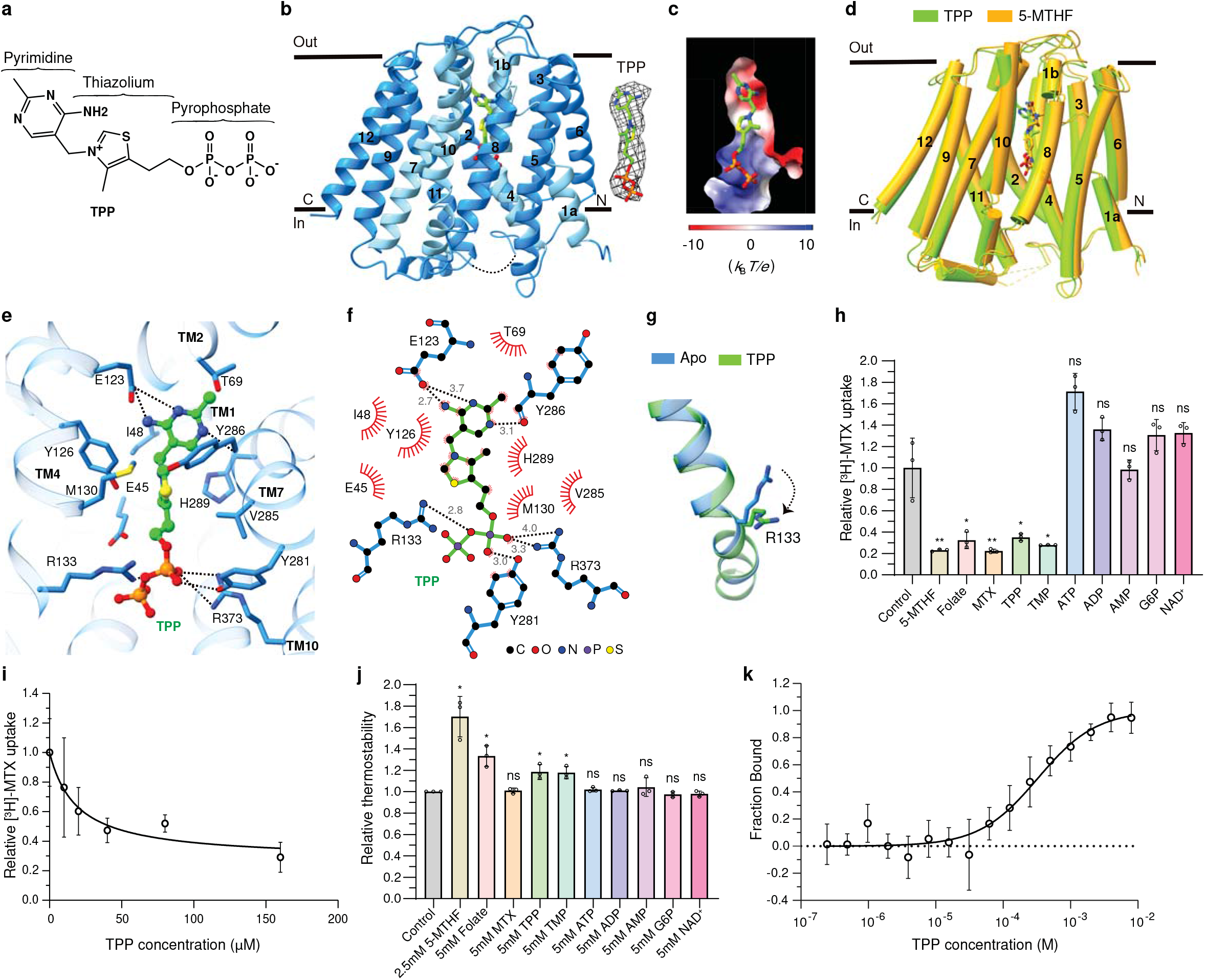
Verification of TPP as the favorite coupled substrate of SLC19A1. **a** The chemical structure of TPP. The constituent moieties are indicated. **b** Ribbon diagram of SLC19A1 in complex with TPP, TPP is shown in sticks and its cryo-EM densities are shown in black meshes. **c** The electrostatic potential of the TPP binding site. **d** Structural comparison of the TPP- and 5-MTHF-bound SLC19A1. **e** Ribbon presentation of the TPP-binding site in SLC19A1. The residues involved in the interaction with TPP are shown with side chains. **f** A schematic summary of the interactions between SLC19A1 and TPP. The distances of the hydrogen bonds are indicated in angstroms. **g** Conformational change of the side chain of Arg133 between apo and TPP-bound SLC19A1 structures. **h** Inhibitory effect of different compounds on the [^3^H]-MTX uptake activity of SLC19A1. All the molecules were tested at the concentration of 200 μM. The results are normalized to the activity of the control experiment in which no inhibitors are added. All experiments were done in triplicates (n = 3, mean ± s.d.). ns, non-significant; * P < 0.01; ** P < 0.005 (T-test). **i** Quantitative measurement of the potency of TPP in inhibiting the [^3^H]-MTX delivery by SLC19A1. The results are normalized to the MTX transport activity of SLC19A1 in the absence of TPP. All experiments were done in triplicates (n = 3, mean ± s.d.). IC_50_ was calculated by fitting to a nonlinear regression model. **j** Quantification of the fluorescence-detection size-exclusion chromatography (FSEC)-based thermostability assay. The concentrations of molecules are indicated. The thermostability is calculated relative to the control experiment in which SLC19A1 was not incubated with any compounds. All experiments were done in triplicates (n = 3, mean ± s.d.). ns, non-significant; * *P* < 0.01 (T-test). **k** MST analysis to examine the affinity of SLC19A1 for TPP. All experiments were done in multiple replicates (n = 6-10, mean ± s.d.).

We then pursued the cryo-EM structure of the SLC19A1/TPP complex to 3.7-Å resolution (Supplementary Fig. S6 and Table S1). Strikingly, TPP could be clearly detected in the same substrate-binding site as in the SLC19A1/5-MTHF structure (Fig. 4b, c). There was no significant conformational difference between the TPP-bound and 5-MTHF-bound SLC19A1, as the RMSD of the two structures was 1.3 Å (Fig. 4d). TPP interacted with SLC19A1 in a manner highly reminiscent of that in 5-MTHF (Fig. 4e, f). On the extracellular side, the pyrimidine ring of TPP formed hydrogen bonds with the side chain of Glu123 and the main-chain carboxyl of Tyr286. In addition, the pyrimidine and thiazolium rings of TPP were stabilized through van der Waals and hydrophobic interactions by the similar collection of residues that interacted with the pterin and *p*-aminobenzoyl groups of 5-MTHF as described above. On the intracellular side, the negatively-charged pyrophosphate moiety of TPP was clamped by the same positively-charged Arg133 and Arg373, as well as the polar interaction with the hydroxyl of Tyr281. Notably, the side chain of Arg133 assumed the unique conformation for a better adaption of TPP (Fig. 4g and Supplementary Fig. S3c).

To validate TPP as an authentic substrate of SLC19A1, we tested its ability to compete with the uptake of [^3^H]-MTX. As the positive control, 5-MTHF, folate, and MTX all effectively blocked the SLC19A1-mediated intake of [^3^H]-MTX at 200 μM concentration. Importantly, TPP exhibited a comparable inhibitory effect with IC_50_ of 19 μM (Fig. 4h, i). In contrast, ATP, ADP, AMP, G6P, and NAD^+^ showed much weaker or no effect on the [^3^H]-MTX transport (Fig. 4h). In parallel, we measured the thermostability of SLC19A1 in the presence of different compounds (Fig. 4j). As expected, the well-documented substrates 5-MTHF and folate significantly enhanced the thermostability of SLC19A1. Among the examined organic phosphate compounds, only TPP could elevate the thermostability of SLC19A1. Furthermore, microscale thermophoresis (MST) analysis confirmed the binding of TPP to SLC19A1 (dissociation constant *K*_d_ = 327 ± 76 μM) (Fig. 4k). All these results together supported that TPP is the favorite coupled substrate of SLC19A1.

## Discussion

SLC19A1 is the first identified folate transporter ubiquitously expressed in tissues and responsible for folate uptake in most types of mammalian cells^9^. Its action is coupled with the counter-transport of organic phosphate anions^40^. Although multiple cellular metabolites such as ATP, ADP, AMP, and NAD^+^ have been documented as the coupled substrates of SLC19A1, we showed with the structural and functional analyses that TPP would be the favorite compared to those commonly recognized ones. According to the alternative access mechanism, SLC19A1 would cycle between the inward-facing and outward-facing conformations to carry its substrates across the cell membrane^41^. In our current structures, the extracellular (5-MTHF) and intracellular (TPP) substrates are bound to the identical site in SLC19A1, similar to that observed in some other antiporters^42,43^. It is plausible that cytosolic TPP could liberate 5-MTHF from the inward-facing SLC19A1 through competition under physiological conditions, and SLC19A would then adopt the outward-facing conformation for releasing TPP and binding extracellular 5-MTHF again (Fig. 5). The complete documentation of such a transport mechanism awaits the future structure of outward-facing SLC19A1. Folate and TPP are the coenzymes generally characterized for anabolism and catabolism, respectively. Therefore, the coordinated exchange of these two molecules by SLC19A1 might represent a novel, intrinsic part of cell metabolic regulation.

**Fig. 5.**
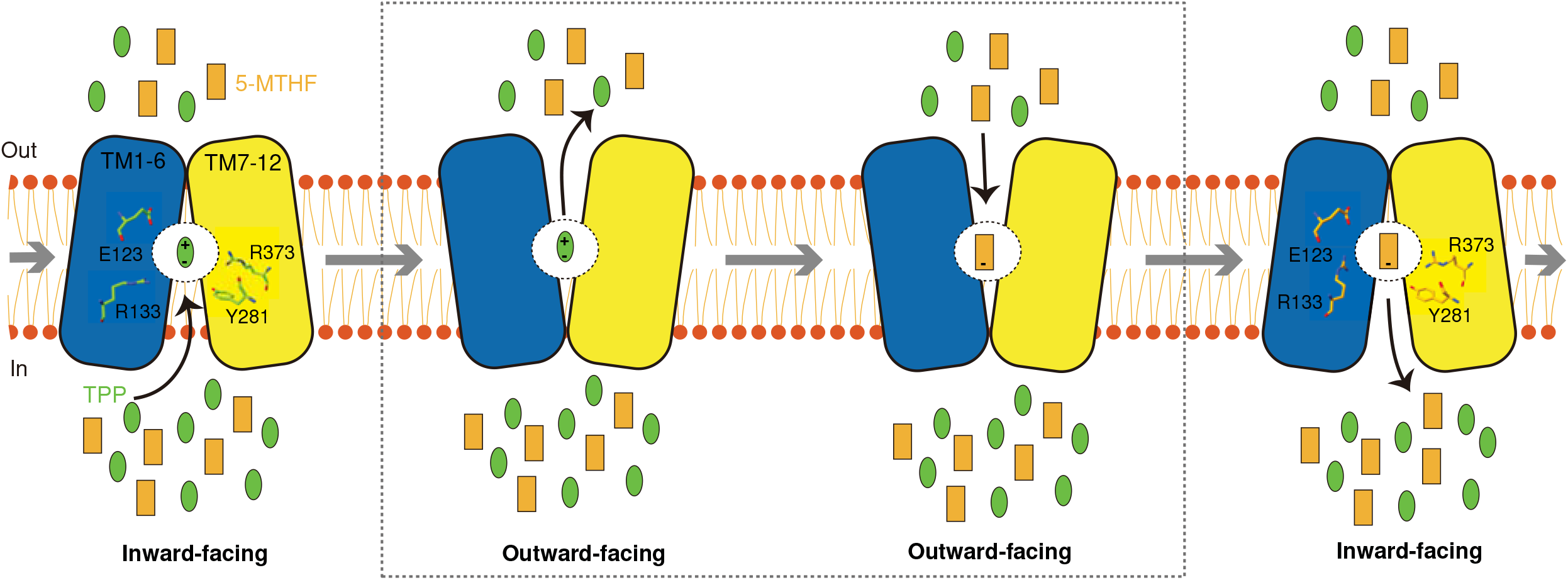
Model of the substrate transport cycle of SLC19A1. SLC19A1 utilizes the alternating access mechanism to reverse transport two substrates. Under physiological conditions, 5-MTHF and TPP are likely the favorite extracellular and intracellular substrates of SLC19A1, respectively. They compete for the same binding site within the central cavity of SLC19A1. The four key residues (Glu123, Arg133, Tyr281, and Arg373) for substrate recognition are indicated.

Thiamine monophosphate (TMP) might bind to SLC19A1 as a coupled substrate. Indeed, TMP effectively competed with [^3^H]-MTX in the transporter assay and enhanced the thermostability of SLC19A1 (Fig. 4h, j). However, given that TMP is an intermediate of thiamine metabolism and its intracellular concentration is approximately one to two orders of magnitude lower than TPP^44,45^, it would likely have a limited role in facilitating folate transport. It is also known that additional organic-phosphate anionic molecules like ZMP (or AICAR, 5-aminoimidazole 4-carboxamide ribonucleoside) and cGAMP (2’3’-cyclic-GMP-AMP) were potential substrates of SLC19A1^46–49^. After the submission of our manuscript, two separate studies reported the structures of SLC19A1 in complexes with some other substrates, including N-hydroxysuccinimide-conjugated MTX (NHS-MTX), pemetrexed (PMX), and different cyclic dinucleotides (CDNs)^50,51^. SLC19A1 exhibited the same inward-facing conformation in all of these structures (Supplementary Fig. S7a). These studies together elucidated the recognition mechanisms for different substrates by SLC19A1. Specifically, folate (5-MTHF) and antifolate drugs (MTX and PMX) bound SLC19A1 as a monomer in a deep cavity very close to the extracellular side, on the contrary, CDNs localized to the broader intracellular entrance of SLC19A1 in the form of a dimer (Supplementary Fig. S7b). Moreover, the interaction mode between SLC19A1 and 5-MTHF was very similar in our and Zhang et al.’s studies, even though the conformations of the pterin ring and glutamate moieties of 5-MTHF showed some differences^51^, indicating these regions might possess some degree of flexibility upon binding to SLC19A1 (Supplementary Fig. S7b, c). However, the antifolate drug MTX was flipped by about 180 degrees along the axis perpendicular to the cell membrane in Wright et al.’s structure^50^, which might be caused by their NHS-mediated crosslinking (Supplementary Fig. S7d).

SLC46A1 and FRs are often specifically expressed or upregulated in cancer cells^52,53^. Of importance, the folate-binding sites of these two proteins are distinct from that of SLC19A1 reported here (Supplementary Fig. S8). Such structural divergence would enable the development of new antifolate drugs that distinguish the different transport systems, thus minimizing potential adverse effects of cancer chemotherapy on normal non-malignant cells. This work has thus bridged a gap in the knowledge of the molecular mechanism of folate transport and could have broad implications for basic biology and translational research.

## Materials and methods

### Cell cultures

HEK293S GnTI^-^ cells were cultured in Yocon HEK293 medium (Yocon Biotechnology) supplemented with 1% FBS and 1% Antibiotic-Antimycotic (Gibco). HEK293F cells were cultured in SMM 293-TI medium (Sino Biological Inc). All the mammalian cells were cultured at 37 °C with 5% CO_2_. Sf9 cells were cultured in Sf-900 II SFM medium (Gibco) at 27 °C.

### Expression and purification of BRIL-SLC19A1

The codon-optimized complementary DNA sequence of BRIL-SLC19A1 was cloned into a pEG BacMam expression vector with 10×His-tag and green fluorescent protein (GFP) attached to the N-terminus^54^. The plasmid was transformed into DH10Bac *Escherichia coli* cells for bacmid generation, and then the bacmid was transfected into sf9 cells using Cellfectin II reagents (Life Technologies) to produce recombinant baculoviruses. For protein expression, 10% of passage 3 (P3) baculoviruses were added to HEK293S GnTI^-^ cells at a density of 3×10^6^ cells/ml. After culturing at 37 °C for 12 hours, 10 mM sodium butyrate was added to boost the protein expression and the cells were transfered to 30 °C. Cells were harvested 48 hours later by centrifugation at 6,000 rpm for 20 min.

For protein purification, the cells were first dispersed by a hand-held homogenizer in lysis buffer (50 mM HEPES pH 7.4, 300 mM NaCl, and 15% glycerol) supplemented with protease inhibitors (1 μg/ml aprotinin, 1 μg/ml leupeptin, 1 μg/ml pepstatin, 20 μg/ml trypsin inhibitor, 1 mM benzamidine, and 1 mM phenylmethylsulfonyl fluoride) and DNase (2 μg/ml), and then lysed by addition of 1% n-dodecyl β-D-maltoside (DDM) and 0.1% Cholesteryl hemisuccinate (CHS) at 4 °C for 2 hours. Cell debris was removed by centrifugation at 18,000 rpm for 40 min. The soluble fraction was mixed with pre-equilibrated anti-GFP nanobody (GFPnb)-coupled cyanogen bromide-activated Sepharose beads (GE Healthcare) at 4 °C for 2 hours. The beads were subsequently washed with 25 column volumes of Buffer A (25 mM HEPES pH 7.5, 150 mM NaCl, and 0.02% DDM-0.002% CHS) and then incubated with PreScission protease (5:1 w/w ratio) at 4 °C overnight to release the target protein. The PreScission protease was removed by incubation with Glutathione Sepharose beads (GE Healthcare) at 4 °C for 1 hour. The protein was further purified by size exclusion chromatography (SEC) using a Superose 6 Increase 10/300 GL column (GE Healthcare) equilibrated with Buffer A. The protein samples of the peak fractions were collected and concentrated to 6 mg/ml using a 100 kDa molecular weight cut-off concentrator (Millipore).

### Expression and purification of anti-BRIL Fab

DNA sequences of the heavy and light chains of anti-BRIL Fab^20^ were cloned into pEG BacMam expression vectors separately. A GFP-tag was attached to the C-terminus of the heavy chain. The bacmids and baculoviruses were prepared in the same way as that of BRIL-SLC19A1. HEK293F cells at a density of 3×10^6^ cells/ml were infected with 10% P3 baculoviruses of heavy and light chains (5% of each). 12 hours after infection, the cells were induced with 10 mM sodium butyrate and transferred to 30°C for protein expression.

For Fab purification, the cell culture media was centrifuged at 6,000 rpm for 30 min, and then the supernatant was concentrated and exchanged into Buffer B (25 mM HEPES pH 7.5 and 150 mM NaCl) using a Hydrosart Ultrafilter system (Sartorius). The following purification steps for anti-BRIL Fab were the same as that of BRIL-SLC19A1. Briefly, after the anti-GFP affinity chromatography, SEC was applied for further purification.

### Expression and purification of anti-Fab Nanobody

The DNA sequence of anti-Fab Nanobody was cloned into pET24 (+) vector which bears an N-terminal PelB signal sequence^55^ and a C-terminal 6×His-tag. The nanobody would be expressed and translocated to the periplasm of *Escherichia coli* strain BL21. The bacteria were cultured in LB medium at 37 °C until OD600 reached about 0.8. Then the protein expression was induced by addition of 1 mM isopropyl-β-D-1-thiogalactopyranoside (IPTG) for 6 hours.

For nanobody purification, the cell pellets were lysed by sonication in lysis buffer. Cell debris was removed by centrifuged at 18,000 rpm for 1 hour. Next, the supernatant was mixed with pre-equilibrated Ni-NTA beads (Smart-Lifesciences) and incubated at 4 °C for 1 hour. The beads were sequentially washed with 20 column volumes of Buffer B containing 25 mM and 50 mM imidazole, and then the target protein was eluted with 3 column volumes of Buffer C (25 mM HEPES pH 7.5, 150 mM NaCl, and 300 mM imidazole). The protein was further purified by SEC using a Superdex 200 Increase 10/300 column (GE Healthcare) equilibrated with Buffer B. The protein samples of the peak fractions were collected and concentrated to 5 mg/ml.

### The BRIL-SLC19A1/Fab/Nb complex assembly

For complex assembly, the purified BRIL-SLC19A1, anti-BRIL Fab, and anti-BRIL Fab nanobody (Nb) were incubated on ice for 1 hour at the ratio of 1:1.4:2. Then the samples were applied to a Superose 6 Increase 10/300 GL column equilibrated with Buffer A to remove the excess Fab and Nb components. The peak fractions corresponding to the ternary complex were collected and concentrated to 6 mg/ml.

### Cryo-EM sample preparation and data collection

Quantifoil R1.2/1.3 Au400 holey carbon grids were glow-discharged for 1 min. 3 μl protein samples were deposited on the grids and blotted for 3 s with filter paper at 10 °C and 100% humidity using a Vitrobot Mark IV (FEI) equipment. The grids were flash frozen in liquid ethane and stored in liquid nitrogen until further use. To prepare the substrate-bound SLC19A1 samples, 5 mM 5-MTHF or TPP was incubated with the BRIL-SLC19A1/Fab/Nb complex on ice for 1 hour prior to grid freezing.

The grids were loaded into a 300 kV Titan Krios electron microscope (FEI) with K3 Summit direct electron detector (Gatan). Data were collected in super-resolution mode at a magnification of ×81,000 using the EPU software (Thermo Fisher Scientific). The physical pixel size was 1.07 Å, and the defocus range was 0.7 to 1.5 μm. The exposure time for each micrograph (40 frames) was 3.2 s, yielding a total dose of 60 e^-^/Å^2^. The numbers of micrographs collected for apo, 5-MTHF-, and TPP-bound SLC19A1 samples were 10,485, 7,382, and 14,262, respectively.

### Cryo-EM data processing

For all the three datasets, the micrographs were motion corrected using MotionCor2^56^ and binned to a pixel size of 2.675 Å. The images with ice contamination were manually removed. Contrast transfer function (CTF) was estimated using Gctf^57^. Particle picking was carried out using Gautomatch (http://www.mrc-lmb.cam.ac.uk/kzhang). Totally 3,104,540 (apo), 1,802,524 (5-MTHF-bound), and 4,144,030 (TPP-bound) particles were automatically picked for the three datasets. After two rounds of 2D classification, the good subclasses were chosen and subjected to the following 3D classification in RELION 3.1^58^. For all the three datasets, one subclass of particles with the highest resolution was re-extracted at the pixel size of 1.07 Å and applied to 3D refinement. The particles and maps of the apo and 5-MTHF-bound datasets were further imported into cryoSPARC for Non-uniform (NU) refinement^59^. The particle numbers used for the final refinement of these three datasets were 268,975 (apo), 206,075 (5-MTHF-bound), and 229,158 (TPP-bound), respectively. The resolutions of the final maps were 3.6 Å for apo SLC19A1, 3.5 Å for 5-MTHF-bound SLC19A1, and 3.7 Å for TPP-bound SLC19A1. All the resolutions reported here were calculated using the 0.143 cutoff criterion. The post-processed maps were generated using DeepEMhancer^60^.

### Model building and refinement

The AlphaFold-predicted model of SLC19A1^61^, and the reported structures of anti-BRIL Fab and anti-BRIL Fab nanobody^20^ (PDB code: 6WW2) were used for our model building. All these models were roughly fitted into the SLC19A1 maps using ChimeraX^62^, and then manually adjusted using Coot^63^. Refinement of the final structures in real space was done by PHENIX^64^. Geometries of the structure models were validated using MolProbity^65^. The Fourier shell correlation (FSC) curves were calculated between the refined models and full maps using PHENIX. Local resolutions were estimated in cryoSPARC. All the figures were prepared using ChimeraX.

### [^3^H]-methotrexate uptake assay

5 μg plasmid of wild-type or mutant SLC19A1 with a C-terminal GFP tag was transiently transfected into 5 ml HEK293F cells at a density of 2-3×10^6^ cells/ml using PEI (BIOHUB). After culturing at 37 °C for 12 hours, 10 mM sodium butyrate was added and the cells were transferred to 30 °C for protein expression. Cells were harvested after 24-30 hours by centrifugation at 4,000 rpm for 5 min. The cells were washed with PBS and resuspended in MHS buffer (20 mM HEPES, and 225 mM sucrose, pH adjusted to 7.4 with MgO) at a concentration of 1.5×10^7^ cells/ml. 10 μl cells were taken out and used for protein expression detection. Next, 100 μl cells were mixed with equal volume of MHS buffer containing 17 nM nonradioactive methotrexate (MTX) and 1.25 pM [^3^H]-methotrexate (0.05 μCi, American Radiolabeled Chemicals, Inc) at 37 °C for 5 min. To analyze the inhibitory effect of TPP, the corresponding molecules were added simultaneously at the indicated concentrations. 800 μl ice cold PBS was added to terminate the MTX transport, and the cells were further washed three times with PBS to remove the extracellular radioactive substrate. Then the cells were lysed with 500 μl 1% Triton X-100 at room temperature for 30 min. 400 μl cell lysate was mixed with 250 μl scintillation fluid, and the [^3^H] radioactive signal was measured using a scintillation counter (PerkinElmer).

To check the protein expression level of SLC19A1, the cells were lysed with RAPI lysis buffer (25 mM Tris-HCl pH 7.5, 150 mM NaCl, 1 mM EDTA, 1% NP-40, and 0.1% SDS) at room temperature for 60 min and mixed with 4×SDS-PAGE loading buffer. The samples were separated using 10 % SDS-PAGE gels. The GFP fluorescence signal of SLC19A1 was directly detected using a gel image system (Tanon).

### Fluorescence-detection size-exclusion chromatography (FSEC)-based thermostability assay

GFP-tagged SLC19A1 expressing cells were lysed with 1% DDM and 0.1% CHS in exactly the same way as that of the BRIL-SLC19A1 purification. After removing the cell debris by centrifugation, the supernatant was incubated with different small molecules on ice for 1 hour. The samples were then treated at 60 °C for 10 min, and subsequently centrifuged at 15,000 rpm for 1 hour to remove the protein aggregates. FSEC was performed using a Shimadzu HPLC system. SLC19A1 protein was separated by a Superose 6 Increase 10/300 GL column during which process the GFP fluorescence signal was monitored. The integral areas of the corresponding fluorescence peak were calculated to represent the amount of SLC19A1 protein. Samples without heat treatment were used as control to evaluate the protein thermostability.

### MST analysis

The affinity of SLC19A1 for TPP was determined using a NanoTemper Monolith NT.115 instrument (NanoTemper Technologies). The full-length SLC19A1 protein was fluorescently labelled with a Red-NHS 2nd Generation kit (NanoTemper Technologies) in Buffer D (25 mM HEPES pH 7.5, 150 mM NaCl, and 0.002% LMNG). 50 nM SLC19A1 was mixed with TPP prepared in 16 different serial concentrations (0.25 μM to 8 mM) in Buffer D. Then the mixture was incubated overnight at 16 °C followed by centrifugation at 15,000 g for 15 min before loaded into the Premium Capillaries (NanoTemper Technologies). The thermophoresis analysis, including six to ten replicate measurements, was performed with 20% excitation power and 60% MST power. The *K*_d_ value was calculated using the MO.Affinity Analysis software (NanoTemper Technologies).

### Data availability

Cryo-EM density maps of the apo, 5-MTHF-bound, and TPP-bound SLC19A1 have been deposited in the Electron Microscopy Data Bank under the accession codes EMD-34817, EMD-34818, and EMD-34819. Their atomic coordinates have been deposited in the Protein Data Bank under accession codes 8HII, 8HIJ, and 8HIK.

## Acknowledgements

We thank the Cryo-EM platform and the School of Life Sciences of PKU for cryo-electron microscopy data collection. We are grateful to Dr. Guopeng Wang, Bo Shao, Xia Pei, and Dr. Ning Gao for their help in cryo-EM experiments. We thank Shitang Huang at the National Center for Protein Sciences of Peking University for assistance with radioactive experiments. We thank Jinkun Xu in Long Li’s lab for assistance with the nanobody screening. We thank Qing Chang and Wenqi Li at Beijing Advanced Innovation Center for Structural Biology of Tsinghua University for assistance with our pilot experiments to verify the protein-small molecule interactions. This research was funded by the National Key Research and Development Program of China (to Z.Z., NO. 2021YFA1302300) and the National Natural Science Foundation of China (to Z.Z., NO. 32171201). The study was also supported by Center for Life Sciences (CLS), School of Life Sciences (SLS) of Peking University, the SLS-Qidong innovation fund, Li Ge-Zhao Ning Life Science Youth Research Foundation, and the State Key Laboratory of Membrane Biology of China.

## Author contributions

Y.D., X.D., and H.Z. did the cloning and protein purification. X.D. screened nanobodies against SLC19A1. Y.D., C.Q., and Z.G. collected the cryo-EM data. Y.D. and Z.Z. processed the EM data. Z.Z. built and refined the structural models. Y.D. and D.Z. performed the MTX uptake experiments. Y.D. carried out the thermostability assay. Y.D. and Y.W. did the MST analysis. C.-H.L. assisted in model building and structural analysis. Z.Z. wrote the manuscript with input and support from all co-authors. J.Y. revised the manuscript.

