## Supplementary Information for "Molecular mechanism of substrate recognition by folate transporter SLC19A1"

**This file includes:**

Supplementary Figures S1 to S8

Supplementary Table S1

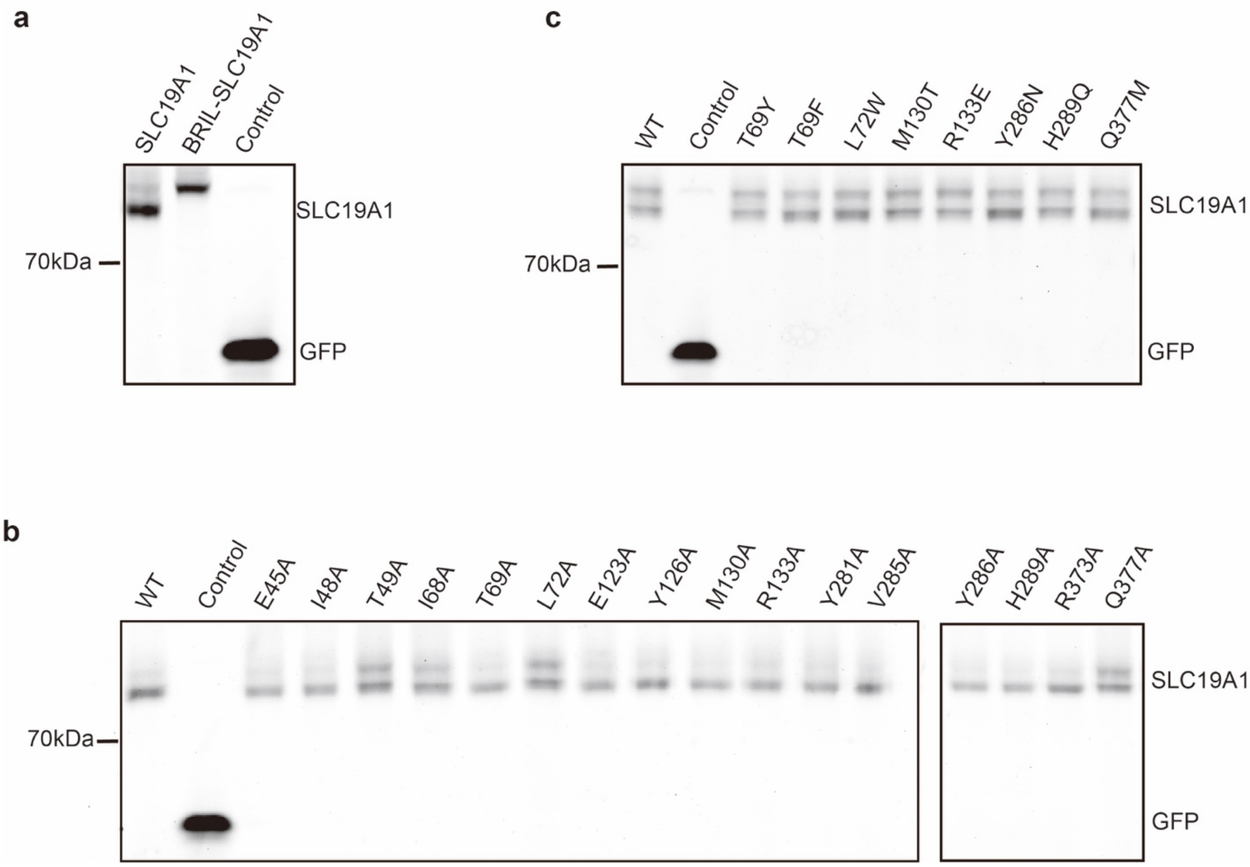


**Supplementary Fig. S1 Detection of the SLC19A1 expression by SDS-PAGE.**

**a-c** Verification of the expression level of different SLC19A1 variants or mutants for the [$^3$H]-MTX uptake assays using the GFP fluorescence signal. All experiments were repeated three times with similar results.

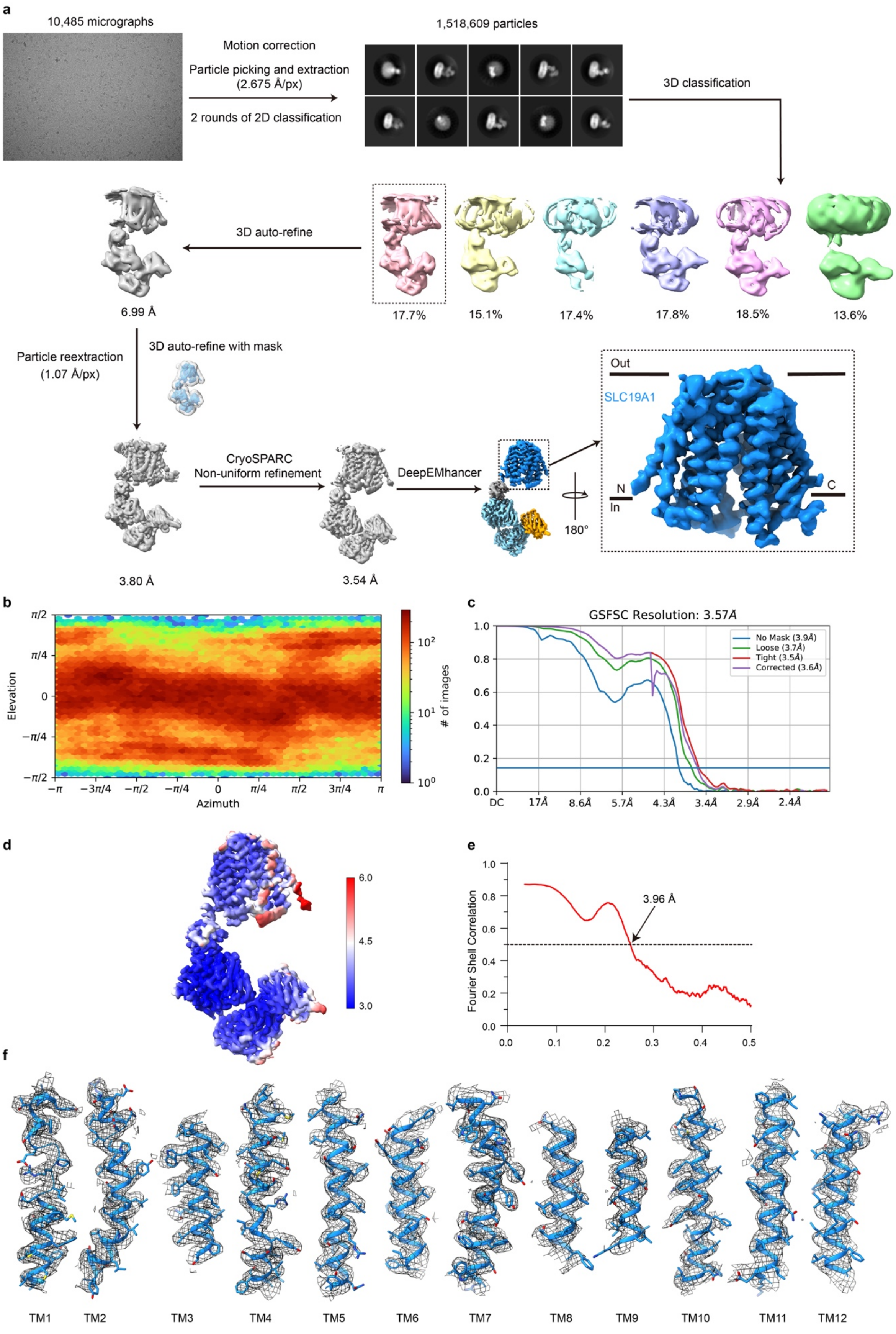
a
10,485 micrographs
Motion correction
Particle picking and extraction (2.675 Å/px)
2 rounds of 2D classification
1,518,609 particles
3D classification
3D auto-refine
6.99 Å
17.7%
15.1%
17.4%
17.8%
18.5%
13.6%
Particle reextraction (1.07 Å/px)
3D auto-refine with mask
CryoSPARC Non-uniform refinement
DeepEMhancer
3.80 Å
3.54 Å
Out
SLC19A1
N
In
C
180°
b
Elevation
Azimuth
### of images
c
GSFSC Resolution: 3.57Å
No Mask (3.9Å)
Loose (3.7Å)
Tight (3.5Å)
Corrected (3.6Å)
DC
17Å
8.6Å
5.7Å
4.3Å
3.4Å
2.9Å
2.4Å
d
6.0
4.5
3.0
e
Fourier Shell Correlation
3.96 Å
f
TM1
TM2
TM3
TM4
TM5
TM6
TM7
TM8
TM9
TM10
TM11
TM12

**Supplementary Fig. S2 Cryo-EM data processing of the apo SLC19A1 dataset.**

**a** Work flow of the data processing. **b** Orientation distribution of the particles used in final reconstruction. **c** Fourier shell correlation (FSC) curves between two half maps. **d** Local resolution of the cryo-EM map. **e** FSC curve calculated between cryo-EM map and structural model. **f** Close-up view of the cryo-EM densities and fitted atomic models for representative regions of the structure.

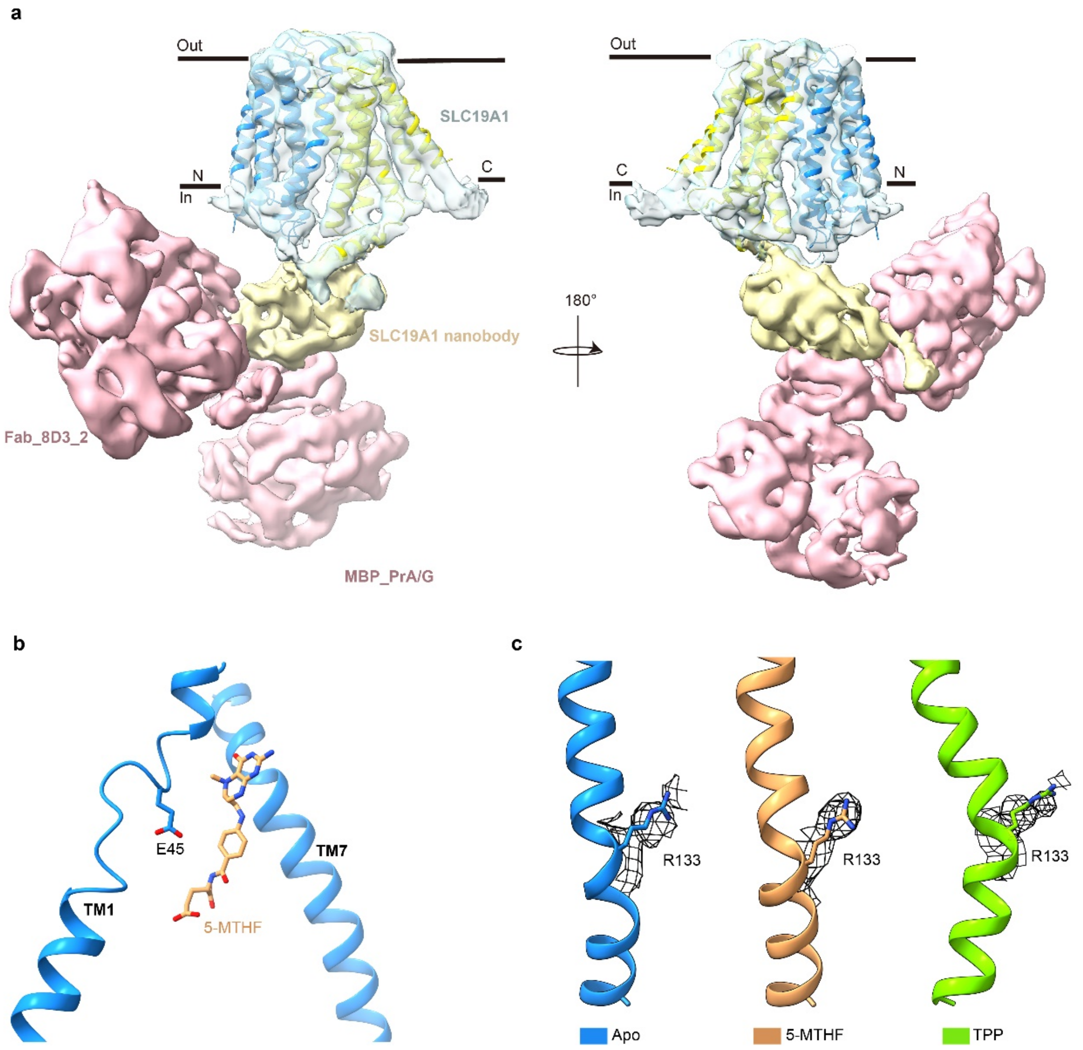


**Supplementary Fig. S3 Structure analyses of SLC19A1.**

**a** Cryo-EM map of the SLC19A1/legobody complex shown in two views. The structure model of SLC19A1 can be fitted well into the EM map. The two half TM bundles of SLC19A1 are colored in blue (TM1-6) and yellow (TM7-12), respectively. The maps of SLC19A1, SLC19A1 nanobody, and the other two components of legobody (MBP_PrA/G and Fab_8D3_2) are colored in light blue, wheat, and pink, respectively. **b** Location of Glu45 in the structure of the SLC19A1/5-MTHF complex. TM1 and TM7 are shown as ribbon. 5-MTHF and the side chain of Glu45 are indicated as sticks. **c** Cryo-EM densities of Arg133 in the three SLC19A1 structures. The side chains of Arg133 are shown as sticks and the cryo-EM densities are shown as black meshes.

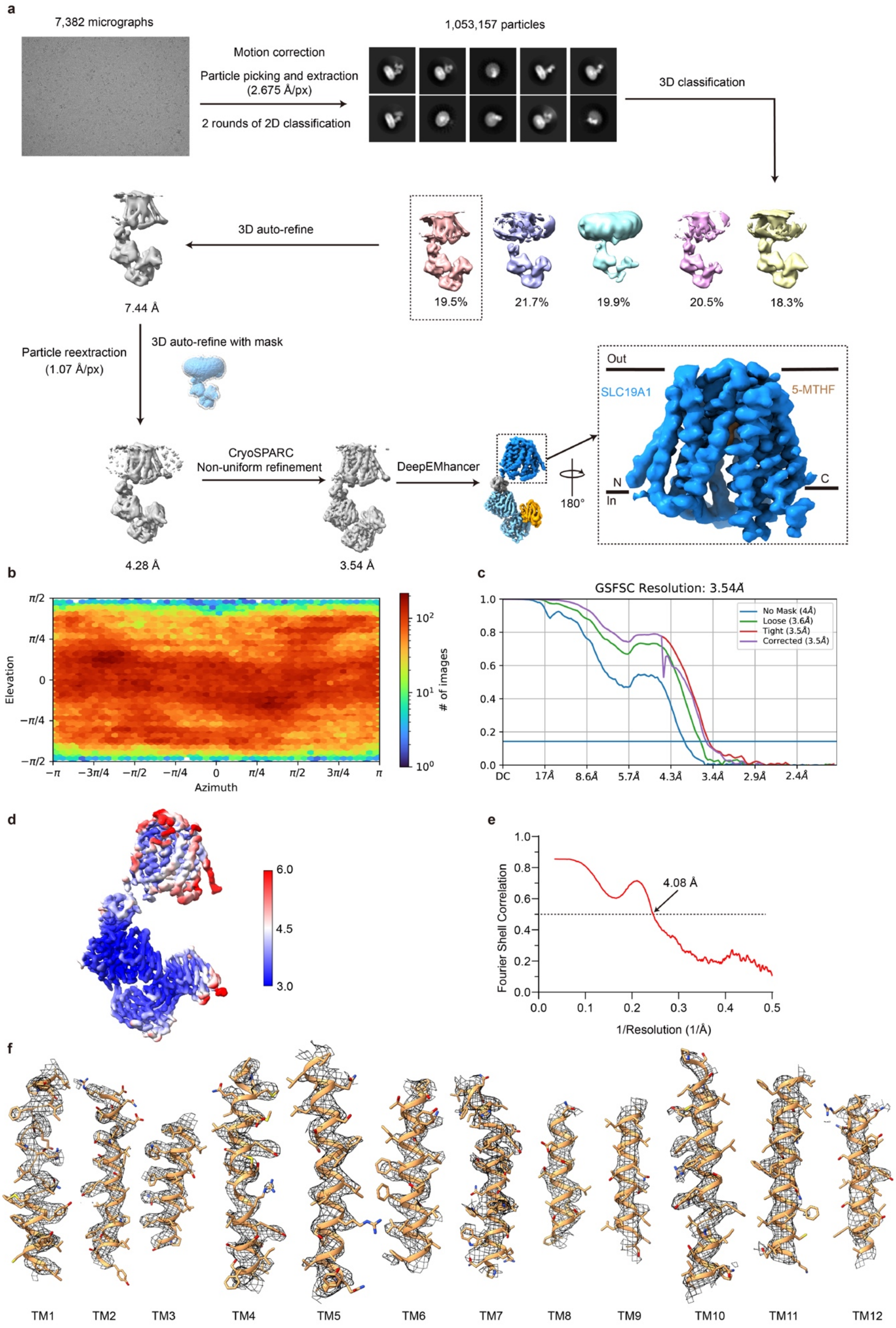
a
7,382 micrographs
Motion correction
Particle picking and extraction
(2.675 Å/px)
2 rounds of 2D classification
1,053,157 particles
3D classification
3D auto-refine
7.44 Å
19.5%
21.7%
19.9%
20.5%
18.3%
Particle reextraction
(1.07 Å/px)
3D auto-refine with mask
CryoSPARC
Non-uniform refinement
DeepEMhancer
4.28 Å
3.54 Å
180°
Out
In
SLC19A1
5-MTHF
N
C
b
Elevation
Azimuth
### of images
c
GSFSC Resolution: 3.54Å
No Mask (4Å)
Loose (3.6Å)
Tight (3.5Å)
Corrected (3.5Å)
DC
17Å
8.6Å
5.7Å
4.3Å
3.4Å
2.9Å
2.4Å
d
6.0
4.5
3.0
e
4.08 Å
Fourier Shell Correlation
1/Resolution (1/Å)
f
TM1
TM2
TM3
TM4
TM5
TM6
TM7
TM8
TM9
TM10
TM11
TM12

**Supplementary Fig. S4 Cryo-EM data processing of the 5-MTHF-bound SLC19A1 dataset.** **a** Work flow of the data processing. **b** Orientation distribution of the particles used in final reconstruction. **c** FSC curves between two half maps. **d** Local resolution of the cryo-EM map. **e** FSC curve calculated between cryo-EM map and structural model. **f** Close-up view of the cryo-EM densities and fitted atomic models for representative regions of the structure.

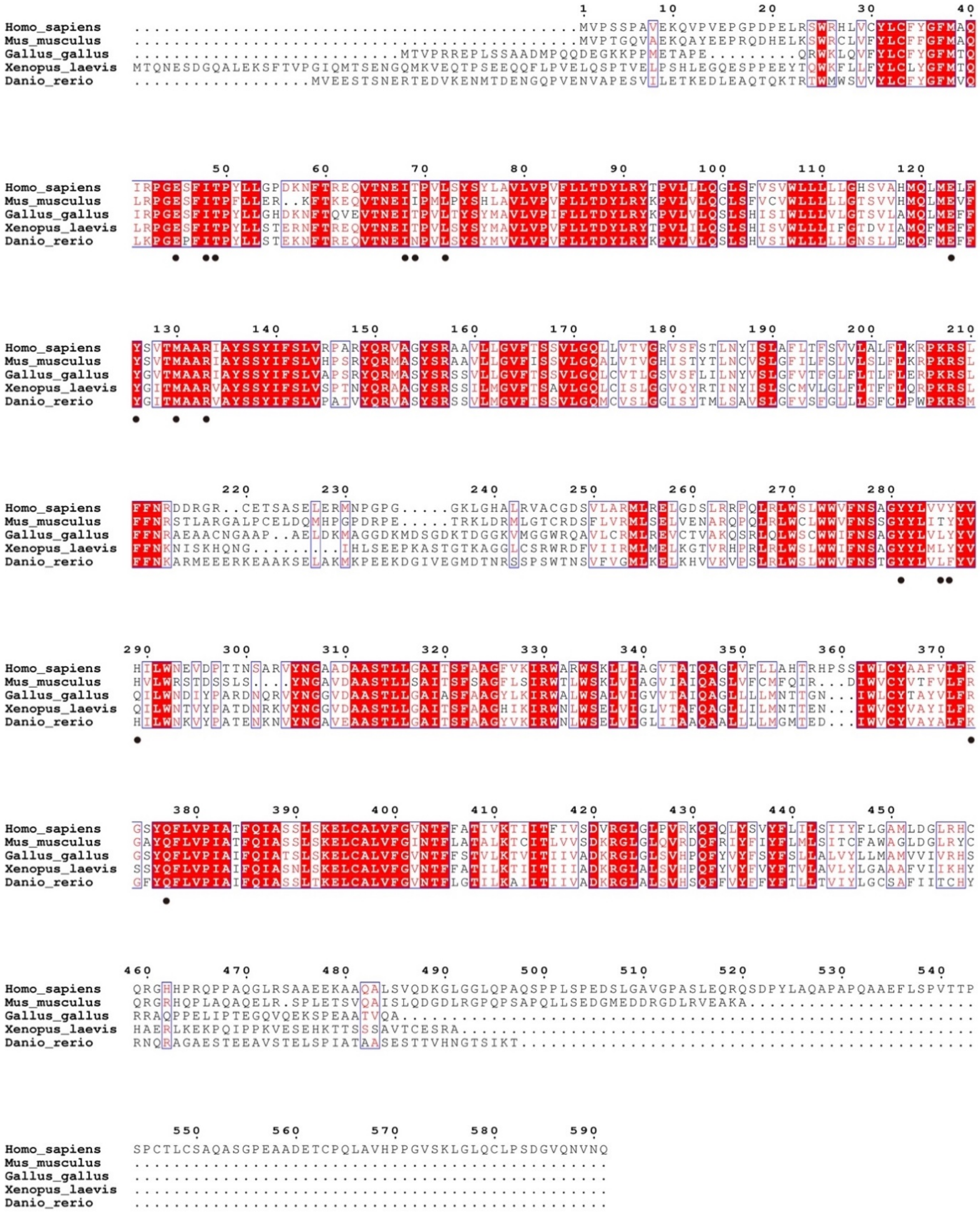

**Supplementary Fig. S5 Sequence alignment of SLC19A1 among several representative species.**

The residues participating into substrate binding are denoted by black dots.

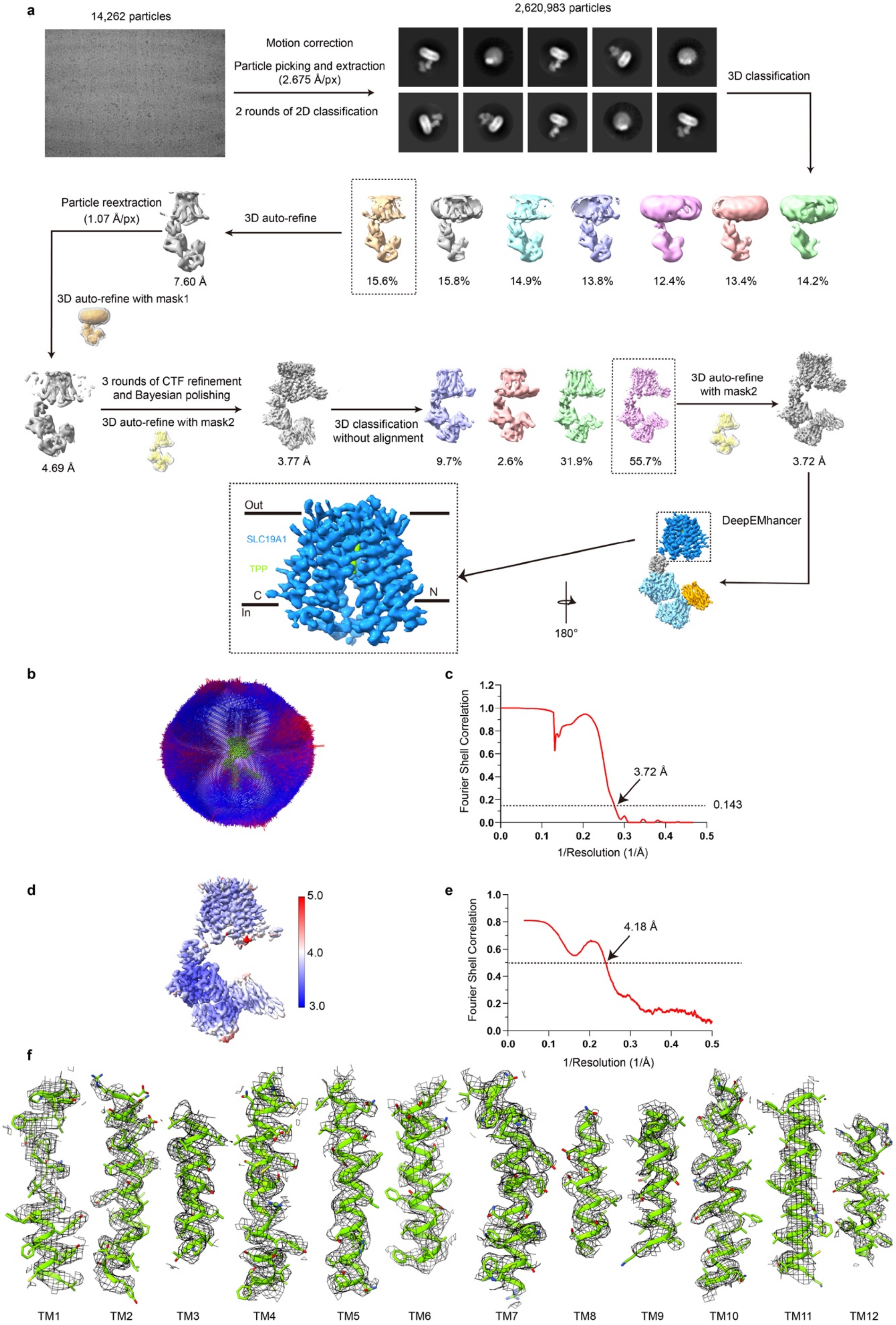
a
14,262 particles
Motion correction
Particle picking and extraction (2.675 Å/px)
2 rounds of 2D classification
2,620,983 particles
3D classification
15.6%
15.8%
14.9%
13.8%
12.4%
13.4%
14.2%
3D auto-refine
Particle reextraction (1.07 Å/px)
7.60 Å
3D auto-refine with mask1
4.69 Å
3 rounds of CTF refinement and Bayesian polishing
3D auto-refine with mask2
3.77 Å
3D classification without alignment
9.7%
2.6%
31.9%
55.7%
3D auto-refine with mask2
3.72 Å
DeepEMhancer
180°
Out
SLC19A1
TPP
C
N
In
b
c
Fourier Shell Correlation
3.72 Å
0.143
1/Resolution (1/Å)
d
5.0
4.0
3.0
e
Fourier Shell Correlation
4.18 Å
1/Resolution (1/Å)
f
TM1
TM2
TM3
TM4
TM5
TM6
TM7
TM8
TM9
TM10
TM11
TM12

**Supplementary Fig. S6 Cryo-EM data processing of the TPP-bound SLC19A1 dataset.**

**a** Work flow of the data processing. **b** Orientation distribution of the particles used in final reconstruction. Red and higher cylinders represent more particles assigned to a particular orientation, while blue and shorter cylinders represent less. **c** FSC curve between two half maps. **d** Local resolution of the cryo-EM map. **e** FSC curve calculated between cryo-EM map and structural model. **f** Close-up view of the cryo-EM densities and fitted atomic models for representative regions of the structure.

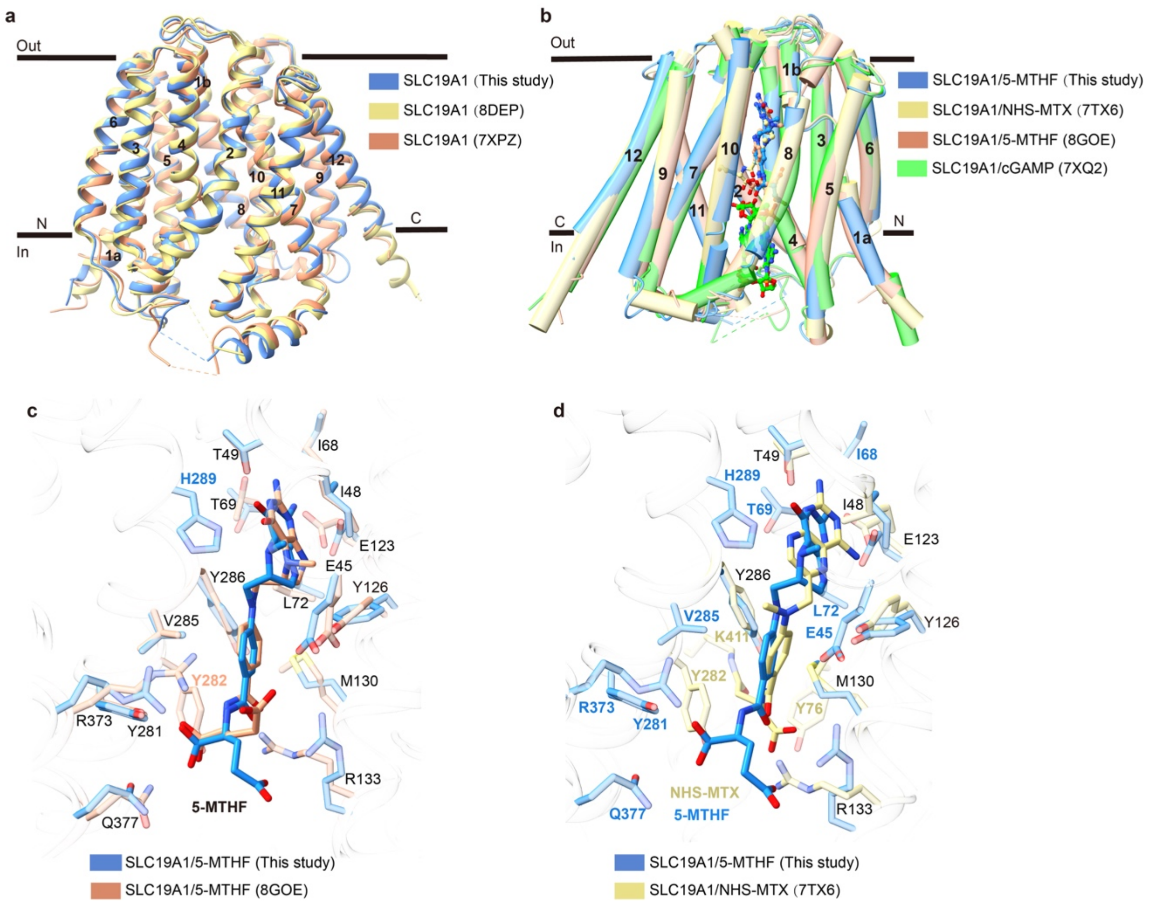


**Supplementary Fig. S7 Superimposition of several SLC19A1 structures.**

**a** Overlay of the apo SLC19A1 structures determined by three separate studies. The RMSDs between these structures are about 0.8 Å. **b** Overlay of the SLC19A1 structures in complexes with different substrates. The RMSDs between these structures are 0.8-1.0 Å. **c** Comparison of the binding details between SLC19A1 and 5-MTHF in our and reported structures. **d** Different conformations of the bound 5-MTHF and NHS-MTX. The residues involved in substrate binding are shown with side chains. Black labels represent the same residues verified in the two structures, while the colored labels represent the unique residues in each structure.

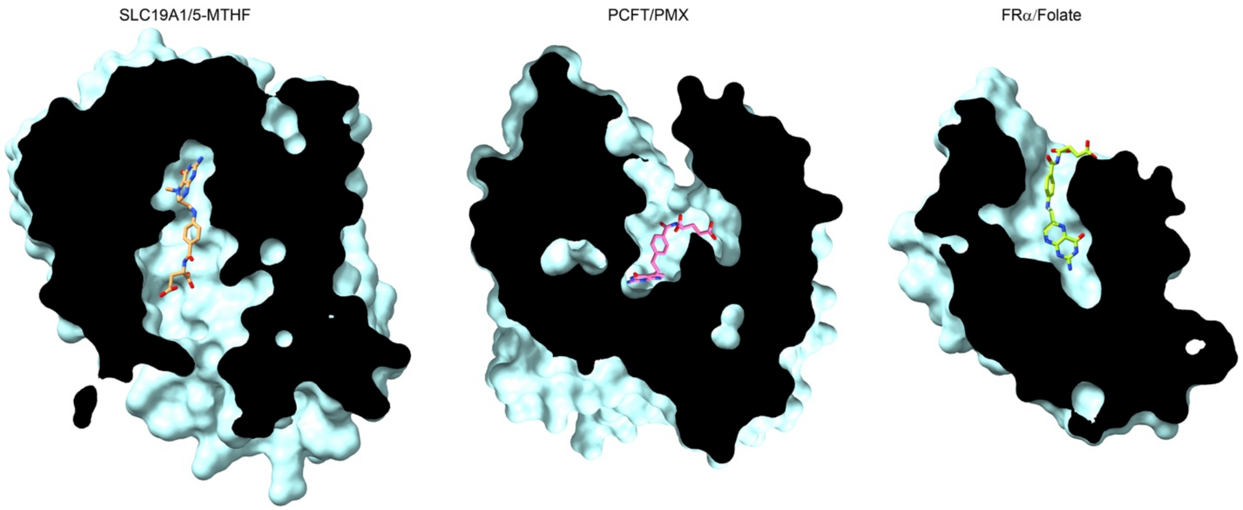


**Supplementary Fig. S8 Structural comparison of the folate binding pockets in SLC19A1 (This study), PCFT (PDB: 7BC7), and FRα (PDB: 4LRH).**

The three proteins are shown as surface presentation. Folate and analogs are shown as sticks.

**Supplementary Table S1 Cryo-EM data collection, refinement, and validation statistics.**

| | Apo (EMD-34817) (PDB 8HII) | 5-MTHF-bound (EMD-34818) (PDB 8HIJ) | TPP-bound (EMD-34819) (PDB 8HIK) |
|---|---|---|---|
| **Data collection and processing** | | | |
| Magnification | 81,000 | 81,000 | 81,000 |
| Voltage (kV) | 300 | 300 | 300 |
| Electron exposure ($e^{-}$/Å$^2$) | 60 | 60 | 60 |
| Defocus range (μm) | 0.7-1.5 | 0.7-1.5 | 0.7-1.5 |
| Pixel size (Å) | 1.07 | 1.07 | 1.07 |
| Symmetry imposed | C1 | C1 | C1 |
| Initial particle images (no.) | 3,104,540 | 1,802,524 | 4,144,030 |
| Final particle images (no.) | 268,975 | 206,075 | 229,158 |
| Map resolution (Å) | 3.57 | 3.54 | 3.72 |
| FSC threshold | 0.143 | 0.143 | 0.143 |
| Map resolution range (Å) | 2.9-12 | 2.9-15 | 3.2-12 |
| **Refinement** | | | |
| Initial model used (PDB code) | 6WW2 | 6WW2 | 6WW2 |
| Model resolution (Å) | 3.96 | 4.08 | 4.18 |
| FSC threshold | 0.5 | 0.5 | 0.5 |
| Model resolution range (Å) | - | - | - |
| Map sharpening *B* factor (Å$^2$) | - | - | - |
| Model composition | | | |
| Non-hydrogen atoms | 8,212 | 8,245 | 8,238 |
| Protein residues | 1,055 | 1,055 | 1,055 |
| Ligands | - | 1 (5-MTHF) | 1 (TPP) |
| *B* factors (Å$^2$) | | | |
| Protein | 66.65 | 96.48 | 70.07 |
| Ligand | - | 80.70 | 67.24 |
| R.m.s. deviations | | | |
| Bond lengths (Å) | 0.003 | 0.003 | 0.003 |
| Bond angles (°) | 0.790 | 0.727 | 0.776 |
| Validation | | | |
| MolProbity score | 1.51 | 1.52 | 1.61 |
| Clashscore | 3.48 | 3.77 | 4.75 |
| Poor rotamers (%) | 0.00 | 0.00 | 0.00 |
| Ramachandran plot | | | |
| Favored (%) | 94.64 | 94.83 | 94.74 |
| Allowed (%) | 5.36 | 5.17 | 5.26 |
| Disallowed (%) | 0.00 | 0.00 | 0.00 |
